# Obligate cross-feeding expands the metabolic niche of bacteria

**DOI:** 10.1101/2020.11.04.368415

**Authors:** Leonardo Oña, Samir Giri, Neele Avermann, Maximilian Kreienbaum, Kai M. Thormann, Christian Kost

**Author notes:** Correspondence (C.K.). Authors contributed equally to this work (Leonardo Oña, Samir Giri, Neele Avermann).

## Abstract

Bacteria frequently engage in obligate metabolic mutualisms with other microorganisms. However, it remains generally unclear how the resulting metabolic dependencies affect the ecological niche space accessible to the whole consortium relative to the niche space available to its constituent individuals. Here we address this issue by systematically cultivating metabolically dependent strains of different bacterial species either individually or as pairwise coculture in a wide range of carbon sources. Our results show that obligate cross-feeding is significantly more likely to expand the metabolic niche space of interacting bacterial populations than to contract it. Moreover, niche expansion occurred predominantly between two specialist taxa and correlated positively with the phylogenetic distance between interaction partners. Together, our results demonstrate that obligate cross-feeding can significantly expand the ecological niche space of interacting bacterial genotypes, thus explaining the widespread occurrence of this type of ecological interaction in natural microbiomes.

## Introduction

Understanding, explaining, and ultimately predicting the distribution of species in space and time is one of the central goals in ecology. The prevailing theoretical framework that is generally applied in this context is based on the niche concept as originally proposed by Grinnell^1^ and Elton^2^, and later on refined by Hutchinson^3, 4^. In his seminal essay^3^, Hutchinson defines the *fundamental niche* as the sum of all abiotic factors that physiologically constrain a species’ performance and survival^4, 5, 6^. The size of the resulting n-dimensional hypervolume is further modified by ecological interactions with other species, thus giving rise to the so-called *realized niche*. Antagonistic interactions for instance, such as predation, parasitism, and competition with other species, can contract the niche space that is available to organisms^5, 7, 8^. From this line of reasoning results the widely-held view that the realized niche is always a subset of the fundamental niche - a notion that is also portrayed in current ecology textbooks^9, 10, 11^. However, ecological interactions can also increase the niche space accessible to a species via a process called *niche construction*. Here, one organism modifies its local environment, thus allowing a second organism to grow and reproduce. As a consequence, the realized niche of a species may actually become larger than its fundamental niche when it synergistically interacts with another species^12, 13^.

In past decades, both theoretical^14, 15, 16, 17^ and empirical research^18, 19^ has primarily focussed on analysing how antagonistic interactions affect the niche space of a given species^20^. However, little is known on whether also synergistic interactions can shape the niche space of interacting individuals^12, 13^. This is surprising given that mutually beneficial interactions are very widespread in natural ecological communities^21, 22, 23, 24^ and have been shown to play key roles in many critical ecosystem processes^23, 25, 26^.

A detailed knowledge of whether and how mutually beneficial interactions deform the ecological niche space of interacting species is crucial for understanding the rules that govern the assembly of complex, multispecies communities^27^. Moreover, the success of attempts to manage natural communities and rationally design functional consortia from the bottom-up rests critically on our ability to predict community structure and function^28, 29^. In addition, when the niche-constructing activities of one species enable another species to invade an environment, in which it otherwise could not survive and reproduce, the resulting interaction is likely to affect the way natural selection operates on one or both species. In this way, an eco-evolutionary feedback loop is closed that can fundamentally alter the ecological role of both species within the community^30, 31^ as well as their prospective evolutionary trajectories^31, 32, 33^. Thus, systematic studies to quantitatively identify the principles that govern niche expansion in synergistic interactions are urgently required.

Here we address this issue by taking advantage of a microbial model system. Specifically, we studied how an obligate exchange of essential amino acids between two bacterial genotypes affects their ability to grow in one of many environments that differed in the carbon source that was available for growth. Comparing the growth of both monocultures in amino acid-supplemented environments to the growth of pairwise consortia in unsupplemented conditions allowed assessing whether the obligate metabolic exchange resulted in an expansion or contraction of the strain’s metabolic niche. For this, we systematically assembled genotypes of five different bacterial species, which were auxotrophic for one of four different amino acids, in all possible pairwise combinations. Analysing their growth as monocultures or pairwise cocultures in one of 31 different carbon environments over time allowed to quantitatively determine the shape of their metabolic niche under the different conditions.

The results of this systematic large-scale experiment revealed first that both niche expansion and niche contraction are common in obligate cross-feeding interactions. Interestingly, niche expansion was significantly more frequent than niche contraction. Second, both the niche space of individual strains and the overlap in niche space of both partners could predict whether the interaction resulted in a contraction or expansion of the niche space occupied by the whole consortium. Finally, the phylogenetic distance between interacting partners was positively associated with niche expansion, yet negatively with niche contraction. Together, our work demonstrates that obligate synergistic interactions frequently expand the ecological niche of interacting partners and identifies some of the key rules determining this effect.

## Results

### Experimental design

To analyse how obligate metabolic cross-feeding affects the ability of bacterial genotypes to grow and survive in different environments, auxotrophic genotypes of five different bacterial species (Table S1) were generated. The resulting mutants were unable to autonomously produce one of four different amino acids (i.e. arginine (*ΔargH*), histidine (*ΔhisD*), leucine *(ΔleuB*), and tryptophan *(ΔtrpB or ΔtrpC*)), thus preventing growth in monoculture without an external supply of the required metabolite. However, coculturing pairs of auxotrophic genotypes can compensate the strains*’* metabolic deficiencies when strains reciprocally exchange essential amino acids^34, 35^. Auxotrophic mutants were grown in monoculture with and without supplementation of the focal amino acid as well as in pairwise cocultures in one of 31 environments, which differed in the carbon source that was available for growth. The total productivity of all communities was quantified in regular intervals (i.e., after 3, 5, and 7 days) by determining cellular respiration (i.e., NADH production).

### Niche expansion is more prevalent than niche contraction

Comparing the expected carbon utilisation profile with the experimentally observed one (Fig. 1) revealed that in 41% ± 1.4% of cases analysed, the growth pattern of monocultures correctly predicted the one observed under coculture conditions (Fig. 2). Strikingly, in 40% ± 2.1% of cases, the obligate synergistic interaction increased the niche space that was available to cocultures, while in 19% ± 0.8% of cases, the extent of niche space was reduced (Fig. 2). Here, the number of carbon sources, cocultures could utilize as a result of niche expansion, was significantly larger than the number of carbon sources, cocultures were unable to metabolize as a result of niche contraction (Kruskal-Wallis test followed by Dunnett’s T3 post hoc test: P = 7.13×10^−8^, χ^2^ = 32.912, n = 324, Fig. 2). Strikingly, 15% of all instances of niche expansion represented cases, in which none of the two auxotrophs could grow in isolation. Thus, in these cases, the ability to use the corresponding carbon sources for growth exclusively emerged from the metabolic interaction among the two auxotrophic genotypes (Fig. S1).

**Fig. 1.**
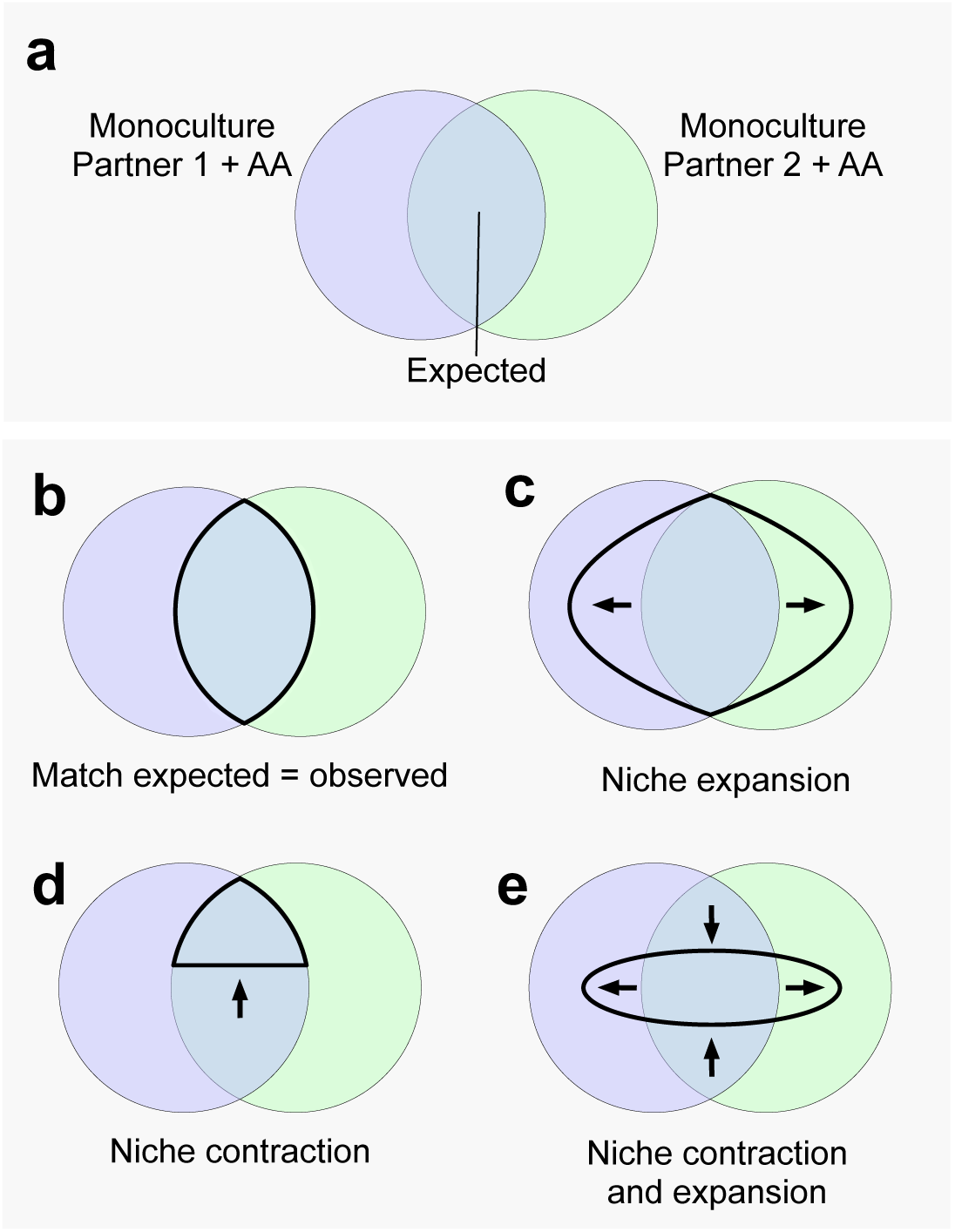
Deformation of niche space in obligate mutualistic interactions. **a** Monocultures of each auxotrophic partner were grown in the presence of the focal amino acid. Circles represent the set of carbon sources, where each partner can grow (i.e., its metabolic niche) and the intersection between sets represents the carbon sources were both partners can grow. **b - e** Observed growth in coculture (thick black set). **b** Perfect match between the expected number of carbon sources used in monoculture and the growth observed in coculture. **c** The niche space occupied by the coculture is larger than the set expected from combined monocultures (i.e., *niche expansion*). **d** The observed set is smaller than the expected set of carbon sources used by both monocultures (i.e., *niche contraction*). **e** Both niche expansion and niche contraction occur simultaneously.

**Fig. 2.**
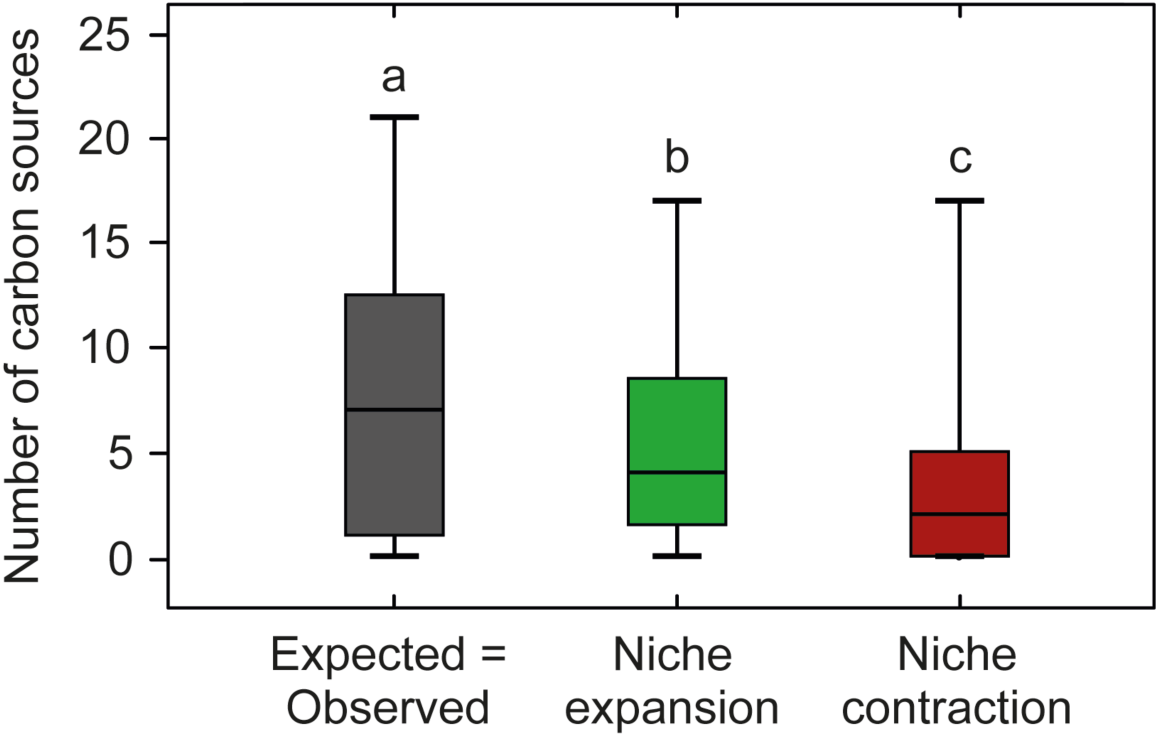
Niche expansion is more common that niche contraction. The overlap in the number of carbon sources used by both the two monocultures (expected) and cocultures (observed) after 5 days of growth was quantified. Comparing the resulting patterns revealed cases, in which the observed niche space matched expectations (grey box) or in which the niche space occupied by cocultures was either larger (green box, *niche expansion*) or smaller (red box, *niche contraction*) than the one predicted by the growth pattern of the two monocultures. Different letters indicate significant differences among groups (Kruskal-Wallis test followed by Dunnett’s T3 post hoc test: P = 7.13×10^−8^, χ^2^ = 32.912, n = 324). Box plots depict median (horizontal line), first and third quartile (box), and the 1.5 inter-quartile range (whiskers).

Additionally, analysing the statistical relation between the degree of niche expansion and niche contraction revealed a negative relationship between both parameters (Spearman rank correlation: P = 5.3×10^−11^, ρ = - 0.58, n = 108, Fig. S2), indicating that the likelihood for niche contraction increases with decreasing niche expansion. These results show that obligate synergistic interactions can strongly affect both size and shape of the ecological niche of two interacting bacterial strains. Notably, the niche space was significantly more often expanded than contracted.

### Metabolic specialisation drives niche deformation

In order to identify the factors that shape the niche space of cocultures, the statistical association between niche expansion or contraction and the size and overlap of monoculture niches was examined. We hypothesized that the ability of genotypes to utilize different carbon sources plays a key role in determining whether a synergistic interaction can successfully establish in a given environment. In this context, the number of carbon sources utilized in monoculture is defined as the *niche width*. Genotypes utilizing a relatively small number of carbon sources have a narrow niche width and are therefore classified as *niche specialists*. Accordingly, genotypes utilizing a larger number of carbon sources have a broad niche width and are thus *niche generalists*.

When the niche width of both partners in monocultures is broad (i.e., when generalists interact with generalists), the probability that partners share carbon sources they both can utilize (i.e., intersection of niche spaces) is expected to be large. Therefore, under this scenario, the number of carbon sources that the coculture can utilize as a result of niche expansion should be small. Cocultures are more likely to expand into niches (i.e., carbon sources) that are utilized by at least one of the two partners. Thus, the potential for niche expansion is expected to be constrained by *niche dissimilarity*, which is given by the sum of carbon sources that is exclusively used by one of the two partners. Conversely, when the niche width of one or both partners in monocultures is narrow (i.e., interactions between specialists), the probability that partners share carbon sources they both can utilize (i.e., *intersection of niche spaces*), is expected to be lower than for the case when generalists interact with generalists. Therefore, in interactions among specialists, the number of carbon sources that the coculture can utilize as a result of niche expansion should be large.

Consistent with these expectations, a Gaussian non-linear model fitted to the dataset identified a peak in niche expansion when specialists interacted with specialists (P < 10^−4^, R^2^ = 0.65, n = 108, Fig. 3a) and a peak in niche contraction, when generalists interacted with generalists (P < 10^−4^, R^2^ = 0.72, n = 108, Fig. 3b). Thus, these results corroborate indeed that the degree of metabolic specialisation of two synergistically interacting individuals determines whether the consortium-level niche is larger or smaller than the niche space occupied by the two interacting individuals in isolation.

**Fig. 3.**
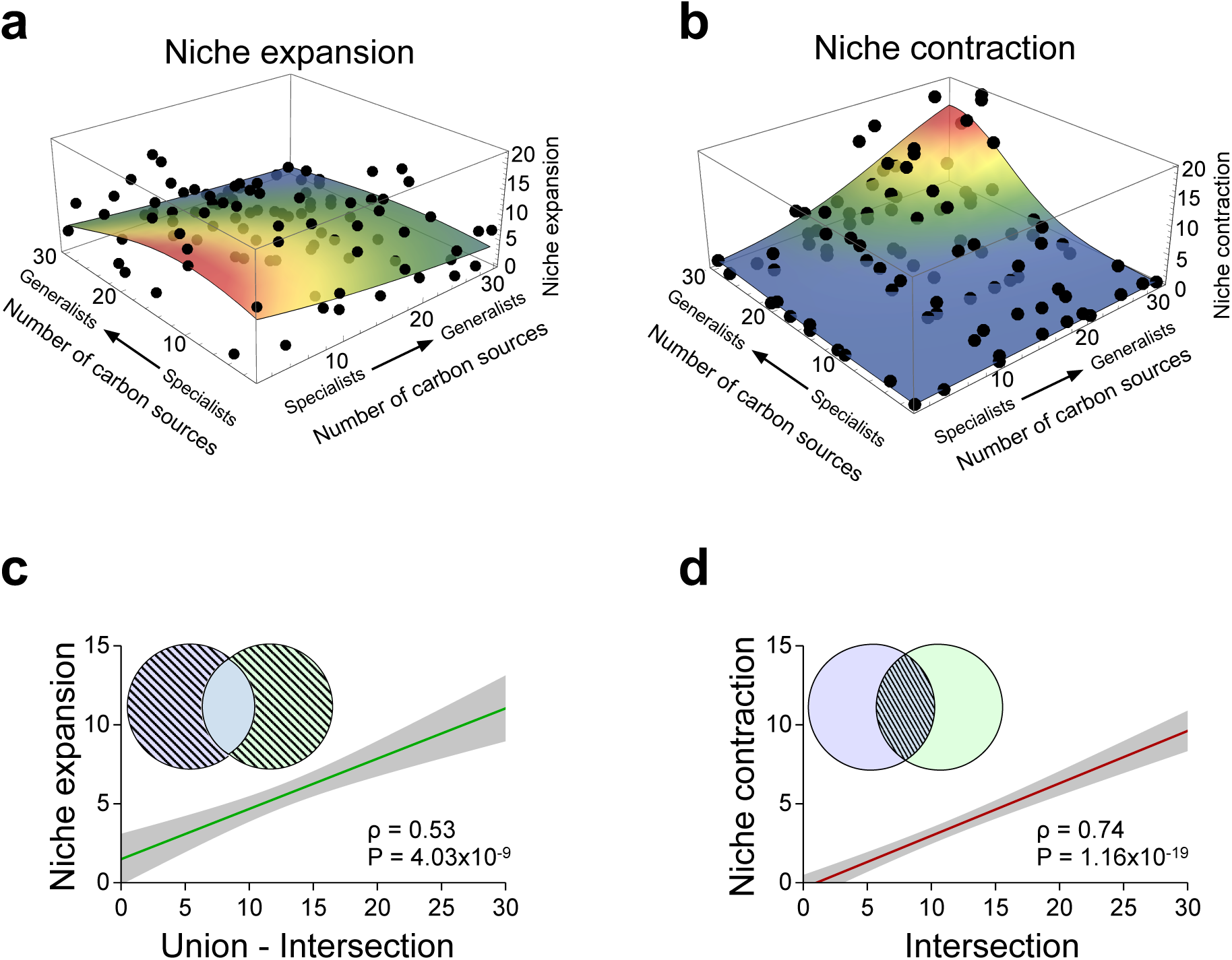
Metabolic specialisation drives niche deformation. The positioning of different bacterial genotypes according to their niche width along the specialist-generalist continuum strongly predicts both niche expansion and niche contraction. **a** Niche expansion was significantly more common when niche specialists interacted with each other (Gaussian non-linear model: P < 10^−4^, R^2^ = 0.56, n = 108), while **b** niche contraction was maximal when generalists interacted with other generalists (Gaussian non-linear model: P < 10^−4^, R^2^ = 0.73, n = 108). **c, d** Monoculture niches and their overlap can predict the magnitude of niche expansion and niche contraction. **c** Niche union minus niche interaction (i.e., *niche dissimilarity*) strongly predicts niche expansion (Spearman rank correlation: P = 4.03×10^−9^, ρ = 0.53, n = 108), while **d** niche overlap (i.e., *niche similarity*) between partners in monocultures strongly predicts niche contraction (Spearman rank correlation: P = 1.16×10^−19^, ρ = 0.74, n = 108). A linear regression was fitted to the data (green/ red line, grey area: ±95% confidence interval).

However, the association of niche expansion and niche contraction with the partners’ niche width does not explicitly consider the degree of similarity between the partner’s niches. A measure of niche dissimilarity is given by the number of carbon sources they do not have in common (i.e., *non-overlap*), which is represented by the union minus the intersection between individual niche spaces (Fig. 3c). Analysing the statistical relation between the resulting value (i.e., *union minus niche intersection*) and the extent of niche expansion revealed a strongly positive correlation between both parameters (Spearman rank correlation: P = 4.03×10^−9^, ρ = 0.53, n = 108, Fig. 3c).

On the other hand, the degree of niche overlap between two monocultures provides a measure of similarity between the two niche spaces and is given by the intersection between them (Fig. 3d). As the potential for niche contraction depends on the number of carbon sources both partners have in common, we hypothesized that niche intersection should be positively associated with the probability for niche contraction. Statistically verifying this hypothesis confirmed that niche overlap (i.e., intersection) between both partners in monocultures was positively associated with niche contraction in coculture (Spearman rank correlation: P = 1.16×10^−19^, ρ = 0.74, n = 108, Fig. 3d).

However, our measures of niche dissimilarity and niche similarity are influenced by the niche width of the interacting partners. For instance, two generalists are expected to have a larger niche intersection than two specialists. To control for this effect, we normalized both niche dissimilarity and similarity with the niche sizes occupied by both corresponding partners (i.e., the niche union). Analysing the linear relationships of both resulting measures with niche expansion and niche contraction corroborated the above findings: the normalized niche dissimilarity was positively associated with niche expansion (Spearman rank correlation: P = 1.5×10^−11^, ρ = 0.59, and n = 108, Fig. S3a) and normalized niche similarity correlated positively with niche contraction (Spearman rank correlation: P = 1.5×10^−15^, ρ = 0.67, and n = 108, Fig. S3b).

Together, these results identify differences in the number and identity of resources used by the interacting genotypes as key determinants of whether an obligate synergistic interaction expands or contracts the metabolic niche of the partners involved. Specifically, metabolic differences between interaction partners favoured niche expansion, while interactions among two more similar types rather resulted in a contraction of the combined metabolic niche.

### Phylogenetic distance between cross-feeding partners predicts niche deformation in coculture

Finally, we asked whether and to which extent niche deformation in coculture was determined by the phylogenetic relatedness among interaction partners (Fig. 4a). Here we anticipated that two closely related species should exhibit a large overlap of their niche space in coculture. This, in turn, should favour a contraction of niche space relative to the two monocultures, because both genotypes are more likely to be similar in resource use and thus compete more strongly. In contrast, the niche space of two more distantly related species is expected to overlap less, thus potentially favouring niche expansion in coculture relative to monoculture conditions. In line with these expectations, a significant negative association was observed between niche overlap and phylogenetic distance (Spearman rank correlation: P = 2.3×10^−11^, ρ = - 0.59, n = 108, Fig. S4). Moreover, niche expansion and phylogenetic distance showed a clear positive association (Spearman rank correlation: P = 5.8×10^−8^, ρ = 0.49, n = 108, Fig. 4b), indicating that the likelihood for niche expansion increases with increasing phylogenetic distance. Furthermore, a significant negative correlation between niche contraction and phylogenetic distance (Spearman rank correlation: P = 1.3×10^−8^, ρ = - 0.51, n = 108, Fig. 4c) implied that the tendency of the combined niche space occupied by a synergistic interaction to contract is increased, when two more related species interact as compared to associations between two more distant relatives.

**Fig. 4.**
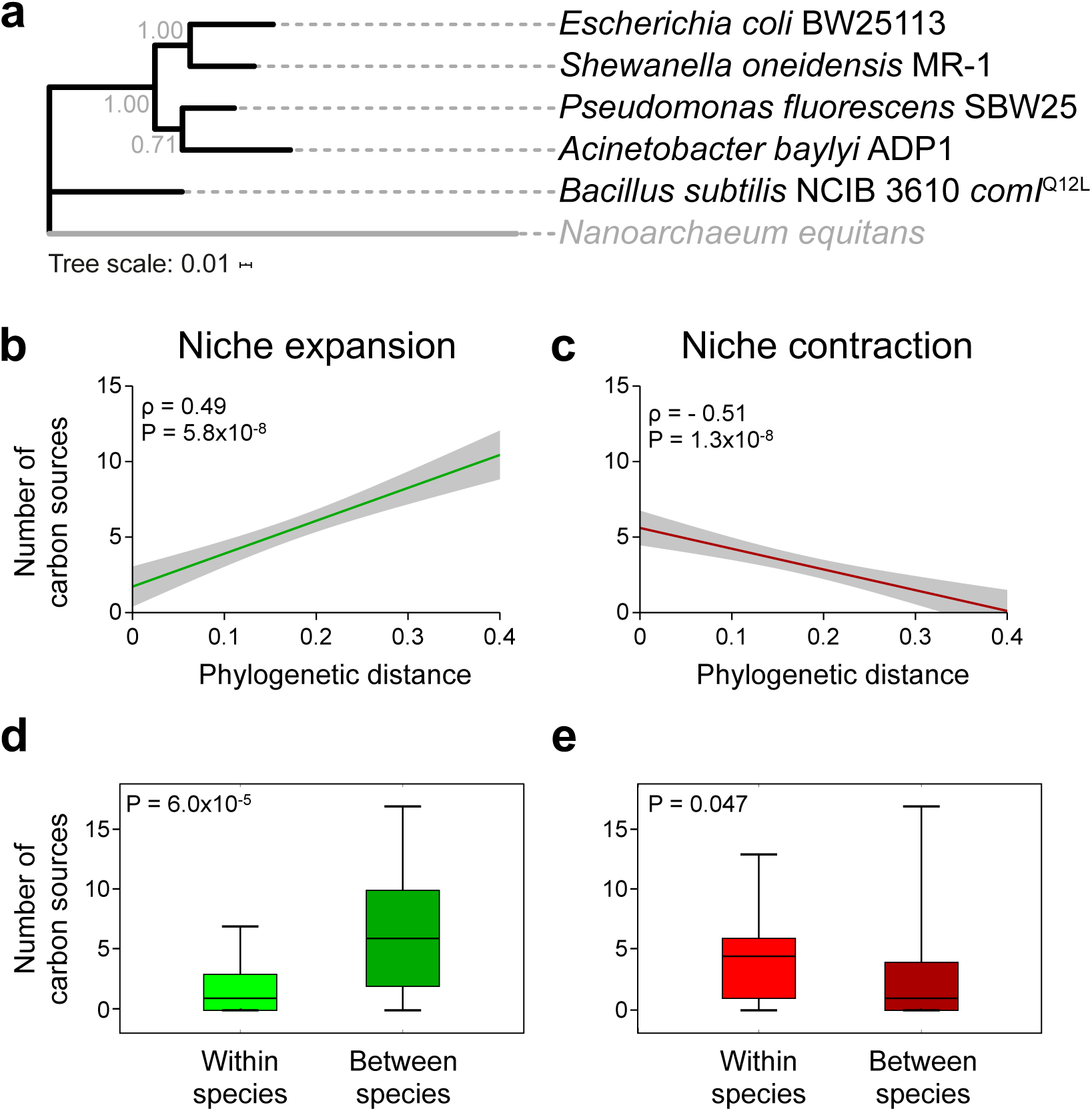
Phylogenetic distance between cross-feeding partners predicts niche deformation in cocultures. **a** Phylogenetic tree of bacterial species used. The outgroup is marked in grey colour and bootstrap values are indicated on nodes. **b-e** Niche expansion and contraction is strongly determined by the phylogenetic relation between both interaction partners. The number of carbon sources corresponds to the number of cases, in which cocultures could or could not grow due to niche expansion and contraction, respectively. **b** Niche expansion is positively correlated with phylogenetic distance (Spearman rank correlation: P = 5.8×10^−8^, ρ = 0.49, n = 108). **c** Niche contraction is negatively correlated with phylogenetic distance (Spearman rank correlation: P = 1.3×10^−8^, ρ = - 0.51, n = 108). A linear regression was fitted to the data (green/ red line, grey area: ±95% confidence interval). **d** Niche expansion was significantly more common when partners belonged to different species (between species) as compared to intraspecific interactions (within species, Mann Whitney U-test: P = 6.0×10^−5^, U = 421.5, n = 108), while **e** the opposite pattern was observed for niche contraction (Mann Whitney U-test: P = 0.047, U = 1202, n = 108). Box plots depict median (horizontal line), first and third quartile (box), and the 1.5 inter-quartile range (whiskers).

Following this logic, niche expansion should be more prevalent in interspecific interactions, while niche contraction should be more common in intraspecific interactions. In line with this hypothesis, we observed that the number of carbon sources cocultures could use as a result of niche expansion was indeed significantly larger for partners belonging to different species as compared to within-species interactions (Mann Whitney U-test: P = 6.0×10^−5^, U = 421.5, n = 108, Fig. 4d). In contrast, interactions between members of the same species were significantly more likely to contract the combined niche space than was the case for interspecific interactions (Mann Whitney test: P = 0.047, U = 1202, n = 108, Fig. 4e).

Taken together, this analysis identified the evolutionary relatedness among two bacterial species as a key predictor of niche expansion or contraction in obligate cross-feeding interactions. In particular, interactions among phylogenetically more dissimilar individuals favoured niche expansion, while an increased relatedness of interaction partners resulted in a contraction of the combined niche.

## Discussion

Identifying the factors that shape the ecological niche of species is a central goal in ecology^6^. While the niche-contracting effect of antagonistic interactions is relatively well understood^19^, our knowledge on how synergistic interactions affect the niche space of species remains poorly developed. Here we identify the phylogenetic distance, niche width, and niche overlap between interacting partners as key factors determining niche deformation in obligate bidirectional cross-feeding interactions between different bacterial genotypes. In systematic experiments, in which hundreds of combinations of paired auxotrophs belonging to different bacterial species were cultivated in a wide range of carbon sources, we show that (i) obligate synergistic interactions are more likely to expand the niche of interacting individuals than to contract it (Fig. 2), (ii) interactions involving niche specialists are more likely to expand the niche space of the interacting consortium, while interactions amongst generalists are more likely to contract it (Fig. 3), and (iii) the phylogenetic distance between strains is positively associated with niche expansion and negatively with niche contraction (Fig. 4).

The amount of resources organisms can allocate to different biological processes is limited. By investing resources in one trait, organisms have to compromise on others^36, 37, 38^. Trade-offs resulting from these evolutionary decisions determine an organism’s niche width and, thus, also its positioning along the specialist-generalist continuum^39, 40^. One major question in ecology is why generalists do not outcompete specialists despite apparent competitive advantages that result from the ability to thrive under a broader range of conditions^41^. Possible explanations of this include the classical antagonistic pleiotropy hypothesis^41^ as well as modern hypotheses that are based on, for example, evolvability costs associated with niche expansion^42^. However, these hypotheses assume a scenario of competition between species and do not take into consideration how mutualistic interactions can affect niche expansion. Our results suggest that when species engage in an obligate, mutually beneficial interaction, one partner in the association can - by facilitation - expand the niche of the respective other one. Our work therefore shows that niche expansion is stronger when partners are metabolic specialists and the overlap between niches is small. Conversely, niche contraction was most prevalent when interactions among niche generalists were considered. This pattern suggests that there is a trade-off between the number of resources that can be exploited and the efficiency of exploitation. A possible explanation for this could be that albeit generalists are able to exploit a wider range of carbon sources, the resulting growth in each one of them might be lower than the growth of a genotype that has specialised to utilize this particular resource. A consequence of this cost of generalism could be that cocultures consisting of generalists are more prone to collapse than cocultures involving specialists. Consequently, generalist genotypes may rather constrain growth of the corresponding consortium, while specialists can efficiently enhance growth of the corresponding partner, thus stabilizing the entire consortium. For a newly established synergistic interaction to prevail in the long-run, the benefits resulting from the interaction have to outweigh the costs. Therefore, the niche-contracting effect of generalist genotypes may represent an opportunity cost with potentially fitness-limiting effects. Future work should analyse whether obligate mutualistic interactions are more likely to evolve from specialist genotypes. Another promising field of research is to understand how the generalist-specialist continuum drives the establishment of metabolic cross-feeding interactions in complex environments.

Identifying the mechanisms that shape the niche space of bacteria (e.g., the number of carbon sources a species can use) and allow different species to coexist, are crucial for developing a quantitative understanding of microbial ecosystems. Given the vital roles of the gut microbiome for human health and disease, the composition and dynamics of mammalian gut microbiota have been subject to intense research in recent years^43, 44^. Processes that structure the gut microbiota are generally believed to be niche-driven, with the spectrum of available nutrients determining whether a microorganism can successfully establish or not^27^. The large taxonomic diversity that is commonly observed in these systems^45^ has been explained as resulting from non-equilibrium conditions that are due to temporal dynamics and spatial heterogeneity, thus preventing competitive exclusion^27^. Even though synergistic interactions are expected to be ubiquitous in these ecosystems as well^46, 47, 48^, little attention has been paid to their role in promoting biological diversity. In fact, recent computational analyses across thousands of environments concluded that host-associated habitats, including gut microbiota, are dominated by highly cooperative communities that are resilient to nutrient change^49^. Our results suggest that the emergence of niche expansion can be a strong ecological force favouring the presence of bacteria in conditions, in which they otherwise would not be able to survive in the absence of their partner. More research is therefore necessary to fully understand how synergistic metabolic interactions expand the niche space of interacting individuals and thus help to stabilize these communities despite fluctuating nutrient availabilities.

Our results revealed that niche expansion is more prevalent among two more distantly related species and less common in intraspecific interactions. Two closely related cells are more likely to share a more similar metabolic network than two distant relatives^35, 50, 51, 52^. As a consequence, the metabolic niche space occupied by two closely related bacteria is more similar than the one of two more distantly related individuals. This relationship can explain why two close relatives are more likely to experience strong competition than the representatives of two different species^19, 53, 54^. In addition, interspecific interactions facilitate metabolic complementarity among interacting partners, thus explaining the observed niche-expanding effect in these cases.

By taking advantage of a synthetic bacterial model system, we systematically investigated in a high-throughput manner how mutually beneficial interactions affect the metabolic niche space of the interacting individuals. Our experimental results demonstrate that synergistic interactions can actually expand the size of the *realized niche* beyond the limits of the *fundamental niche*, an observation is in stark contrast to classical niche theory^7, 8, 55^. Experiments, such as the one presented in this work, can help to uncover the basic principles governing the emergence and maintenance of diversity in complex microbial ecosystems. Moreover, ecological niche-based predictions are an underused, but highly promising tool to guide the design of rationally engineered microbiomes for medical or biotechnological applications. Future studies using natural and laboratory-based systems should analyse how other, more complex interactions determine the niche space of the participating individuals and how these changes create eco-evolutionary feedbacks that in turn might affect fundamental process such as ecosystem stability and function.

## MATERIALS AND METHODS

### Bacterial strains

The following five, bacterial species were used: *Acinetobacter baylyi* ADP1, *Bacillus subtilis* 3610 *comI*^Q12L^, *Escherichia coli* BW25113, *Pseudomonas fluorescens* SBW25, and *Shewanella oneidensis* MR-1 (Table S1). This included both the prototrophic wild type of each species as well as strains that were auxotrophic for one of four amino acids arginine (*ΔargH)*, histidine (*ΔhisD)*, leucine *(ΔleuB*), and tryptophan *(ΔtrpB* or *ΔtrpC)* (Table S1).

### General cultivation conditions

All strains were cultured at 30 °C in modified minimal medium for *Azospirillum brasiliense* (MMAB)^56^ with N-acetyl-glucosamine (NAG) (5 gL^-1^) as a sole carbon source, where all species showed considerable growth. For strain constructions, strains were grown in lysogeny broth (LB) or on LB agar plates at 30 °C. MMAB medium consists of; 3 gL^-1^ K_2_HPO_4_, 1 gL^-1^ NaH_2_PO_4_, 0.15 gL^-1^ KCl, 1 gL^-1^ NH_4_Cl, 5 mL^-1^ from 60 gL^-1^ solution MgSO_4_·7H_2_O, 0.5 mL^-1^ from 20 gL^-1^ solution CaCl_2_·2H_2_O, 0.25 mL^-1^ of 0.631 g 50 ml^-1^ solution FeSO_4_, and trace salts 10 mL^-1^. Composition of 1L trace salts solution: 84 mg L^-1^ of ZnSO_4_·7H_2_O, 765 μL from 0.1 M stock of CuCl_2_·2H_2_O, 8.1 μL from 1 M stock of MnCl_2_, 210 μL from 0.2 M stock of CoCl_2_·6H_2_O, 1.6 ml from 0.1 M stock of H_3_BO_3_, 1 ml from 1.5 g/100 ml stock of NiCl_2_. LB medium consists of 10 gL^-1^ tryptone, 5 gL^-1^ yeast extract, 5 gL^-1^ NaCl. For solidification, agar was added to a final concentration of 1.5% (wt/vol%). For the generation of auxotrophic strains, antibiotics were used at the following final concentrations: kanamycin 50 µg ml^-1^ (*E. coli* and *A. baylyi*), nitrofurantoin 100 µg ml^-1^, tetracycline 15 µg ml^-1^ (*P. fluorescens*), and erythromycin-lincomycin (2 µg ml^-1^ erythromycin + 12.5 µg ml^-1^ lincomycin; *B. subtilis*). To generate auxotrophic strains of *P. fluorescens*, X-gal (5-bromo-4-chloro-3-indolyl-β-D-galactopyranoside) was added to agar plates at a concentration of 50 µg ml^-1^. Addition of diaminopimelic acid (DAP, final concentration: 300 μM) allowed growth of the conjugation strain *E. coli* WM3064 (Table S2).

All genotypes were streaked to a single colony to confirm their purity. To prepare precultures, glycerol stocks frozen at −80 °C were streaked on LB agar plates and grown for 24 h at 30 °C. All optical density (OD) measurements were performed with a FilterMax F5 microplate reader (Molecular Devices) at 600 nm. In all the experiments, independent biological replicates were used.

### Construction of auxotrophic strains

*Escherichia coli* BW25113 and *Acinetobacter baylyi* ADP1 wild type were used as parental strain to construct auxotrophic mutants. In both species, target genes were replaced with a kanamycin resistance cassette. In *E. coli*, auxotrophs were generated via P1 phage transduction^57^ and in *A. baylyi* as described^58^. In addition, antibiotic resistance was cured by excising the kanamycin cassette from the genome of the auxotrophs of both species. For this, the pCP20 plasmid was used, which harbours the FLP recombinase^59^.

#### Construction of *Bacillus subtilis* 3610 comI^Q12L^ auxotrophs

All *B. subtilis* strains generated in this work were obtained by using natural competence transformation with genomic DNA of donor strains. *Bacillus subtilis* 3610 *comI*^Q12L^ (DK1042 is the naturally competent version of NCIB 3610) was used as both wild type strain and for construction of auxotrophic mutants^60^. Cells were grown in LB broth medium at 37 °C with aeration or on LB agar plates. Auxotrophic strains were obtained by naturally transforming genomic DNA from *B. subtilis* 168 (Δ*argH*, Δ*hisD*, Δ*trpC*, and Δ*leuB*) genotypes, respectively^61^ (Table S1). Transformation was achieved in the following way: wild type *Bacillus subtilis* 3610 cells were streaked out on agar plates and incubated at 37 °C. The next day, a colony was picked, inoculated into 2 ml of complete competence medium (MC 10x for 100 ml; 14.0 g K_2_HPO_4_, 5.2 g KH_2_PO_4_, 20 g glucose, 8.8 g trisodium citrate dihydrate, 2.2 g ferric ammonium citrate, 1 g casamino acids, 2 g potassium glutamate monohydrate, for 1x added 1.8 ml H_2_O ml, 200 µl of 10x MC and 6.7 µl 1M MgSO_4_), and grown at 37 °C for five hours. 400 µl of the resulting culture was then mixed with genomic DNA in a fresh 15 ml tube and incubated for 2 h at 37 °C. Afterwards, cells were plated on LB agar containing the selective antibiotic erythromycin (1 µg ml^-1^ erythromycin + 12.5 µg ml^-1^ lincomycin). After 24 h of incubation, 8 different single colonies from each plate were purified by re-streaking on a fresh selection plate. As the genomic DNA from the donor *B. subtilis* 168 contained a gene encoding erythromycin (Erm^R^) resistance, colonies were selected using erythromycin. After overnight incubation, six individual colonies were purified and stored in 50 % glycerol at −80 °C.

To eliminate the antibiotic resistance cassette from the genome, auxotrophs were transformed with the pDR244 temperature-sensitive plasmid, which constitutively expresses the Cre recombinase and is marked with a spectinomycin resistance cassette^61^. Transformants were selected on LB supplemented with 100 μg ml^-1^ spectinomycin at 30 °C - a permissive temperature for pDR244 replication. Transformants were then streaked on LB without antibiotic and incubated at 42 °C - a non-permissive temperature for plasmid replication. Single colonies were re-streaked on LB, LB with spectinomycin, and LB with erythromycin-lincomycin, and everything was incubated at 37 °C. Isolated clones that could grow on LB, but were sensitive to the two antibiotics had lost pDR244 and the resistance cassette. *Cre/lox* mediated loop-outs of the antibiotic resistance cassette from auxotrophic strains were confirmed by PCR with oligonucleotide primers flanking the deletion^61^ (Table S3).

#### Construction of *Pseudomonas fluorescens* SBW25 auxotrophs

Auxotrophic genotypes (Δ*argH*, Δ*hisD*, Δ*trpB*, and Δ*leuB*) were constructed using a scar-free, pUIC3-mediated, two-step allelic exchange protocol of *P. fluorescens* SBW25^62, 63^. Primers used are indicated in Table S3. Gene deletions were obtained by splicing via overlapping extension PCR (SOE-PCR) with a two-step allelic-exchange strategy using the suicide-integration vector pUIC3 as previously described^63, 64^. PCR was used to amplify an approximately 1 kbp upstream and downstream region surrounding each auxotrophy-causing mutation and the product was cloned into pCR8/GW/TOPO plasmid using the TA cloning kit (Invitrogen). Sequence identity was verified by DNA sequencing and the correct DNA fragment was then cloned into the *Bgl*II site of pUIC3 vector^63^. Allelic exchange and the construction of pUIC3 plasmid was performed using *E. coli* DH5α λpir. The resulting pUIC3 construct was then conjugated into SBW25 with the help of plasmid pRK2013 using the standard protocol^63, 65^, in which the cloned fragment integrates into the chromosome by homologous recombination (i.e., first recombination event). Double crossover mutants were selected using cycloserine enrichment^66^. To achieve the genomic replacement (i.e., second recombination event), 10 µl of the transconjugant *P. fluorescens* overnight culture was inoculated into 200 ml liquid LB medium in a 1 l flask. No antibiotics were added to the medium in order to allow loss of the chromosomal pUIC3 construct. Following 16 h of incubation at 28 °C (150 rpm), 100 µl was transferred to 20 ml fresh LB in a 250 ml flask, and incubated for 30 min at 28 °C at 150 rpm. After that, 10 µg ml^-1^ tetracycline was added prior an additional 2 h-incubation step at 28 °C, in order to select for cells, in which the pUIC3 vector has been integrated into the host chromosome. After 2 h, cycloserine (1 g ml^-1^) enrichment was used to kill growing *Pseudomonas* cells and the flask was incubated for 5 h at 28 °C. By killing the growing tetracycline-resistant cells, addition of cycloserine selected for cells that had undergone the rare second round of homologous recombination leading to a loss of the chromosomally inserted vector. Cells were then harvested by centrifugation, washed with LB, and suitable dilutions were plated onto LB agar plates containing X-gal, which was followed by an incubation at 28 °C for 48 h. To screen for a loss of *lacZ*-encoding pUIC3, white colonies were picked and purified by re-streaking and cross checking their tetracycline sensitivity. The region containing the desired marker-less auxotrophic mutation was confirmed by PCR with (UF and DR) primers flanking the deletion (Table S3).

#### Construction of *Shewanella oneidensis* MR-1 auxotrophs

Marker-less, in-frame deletion mutants of *S. oneidensis* MR-1 were generated by sequential homologous cross-over using the suicide vector pNTPS138-R6K as described previously^67^. To this end, up- and downstream fragments of about 500 bp of the focal genes (Δ*argH*, Δ*hisD*, Δ*trpB*, and Δ*leuB*) were amplified by PCR using the corresponding forward and reverse primers (Table S3). The resulting fragments were joined and cloned into EcoRV-linearized vector by Gibson assembly^68^. The resulting plasmid was introduced into *S. oneidensis* MR-1 by conjugation using *E. coli* WM3064 as a donor.

To determine conditional lethality of all engineered auxotrophs, strains were grown in MMAB minimal medium with NAG (5 gL^-1^) as carbon source, which was either supplemented with the focal amino acid (150 µM) to complement the respective mutation (arginine (Δ*argH*), histidine (Δ*hisD*), tryptophan (Δ*trpB*) and leucine (Δ*leuB*)), or left unsupplemented. In all engineered auxotrophic genotypes except *Bacillus subtilis* Δ*trpC* and *Pseudomonas fluorescens* Δ*argH* and Δ*hisD*, the prototrophic phenotype was successfully restored by supplementation of the focal amino acid, while strains were unable to grow in its absence. This confirmed the desired auxotrophic phenotype in the generated strains. The three genotypes that did not show the expected phenotypes were excluded from the main experiment. In total, 17 auxotrophic genotypes of five different bacterial species were used (Table S1).

### Phylogenetic distance

To generate the phylogenetic tree of five different species, the 16S rRNA gene sequences (∼1.5 kbp) of all five bacterial species and one outgroup were retrieved from NCBI GeneBank (Table S1). The tree was constructed using maximum likelihood method in MEGA X software^69^, selecting the Kimura 2-parameter model. Bootstrapping was carried out with 1,000 replicates. The resulting phylogenetic tree was edited using iTOl^70^ and the phylogenetic distances between species were derived from the resulting distance matrix (Table S4).

### Phenotypic growth profiling via single carbon source utilization

The *niche structure* (i.e., the pattern and types of carbon sources used) and *niche width* (i.e., the number of carbon sources used) were evaluated by quantifying the catabolic activities of i) cocultures of auxotrophic strains, ii) monocultures of auxotrophic strains, iii) monocultures of auxotrophic strains, to which the focal amino acid has been added (150 µM), and iv) monocultures of the prototrophic wild types of all species. Carbon source utilization of each strain alone or in coculture was probed using the BiOLOG EcoPlate (BiOLOG, Hayward, CA, USA), which contains 31 different carbon sources in three replicates. The BiOLOG assay uses the tetrazolium violet redox dye to monitor cell respiration. If the focal microorganism utilizes the respective carbon source, oxidation of the nutrient will lead to respiration, which in turn will change the colour of the dye from colourless to purple^71^. Water without any carbon source served as control in triplicates.

Plates were inoculated as follows. First, strains were precultured in replicates by picking single colonies (8 replicates per strain) from LB agar plates and incubated for 24 h in 2.3 ml of MMAB medium + N-Acetyl-Glucosamine (NAG) + 2% LB at 30 °C. Growth of precultures was verified by quantifying their optical density (OD_600nm_) in a plate reader. Four independent replicates that showed approximately the same growth were combined in one 15 ml falcon tube and the OD_600nm_ of these mixtures were measured again. Precultures were then diluted to an OD_600nm_ of 0.05 for all strains by adding MMAB medium without carbon source. To prepare the culture to be transferred into the BiOLOG EcoPlate, approximately 100 μl of precultures were inoculated into 900 μl of MMAB without carbon source. In the case of cocultures, two auxotrophs were mixed in a 1:1 ratio by coinoculating 50 μl of each diluted preculture. To cultivate monoculture with amino acid, 150 μM of the respective amino acid were added. All cultures had a starting density of 0.005 OD_600 nm_. Finally, each well of the BiOLOG EcoPlate was inoculated by 140 μl of culture and plates were sealed with a transparent film. To determine the growth and carbon source utilization pattern, measurements were recorded at four time points: directly after inoculating 0 (i.e., t_0_), after 3 (t_3_), 5 (t_5_), and 7 (t_7_) days, and absorbance at 595 nm was measured. All treatments (cocultures without amino acids) and control (wild types and auxotrophs with/ without focal amino acid) groups used in this experiment were independently replicated three times, resulting in 147 cultures in total. All BiOLOG plates were incubated for 7 days at 30 °C and shaken at 150 rpm (Table S2).

### Data processing and statistical analysis

#### Phenotypic profiles and mathematical definitions of ecological niche properties

To analyse the data resulting from the large-scale cultivation experiment, negative measurements were set to 0 in order to avoid mathematical errors when subtracting values from each other. The absorbance of the water control was subtracted for each well to correct for basal growth in all treatments and controls. Afterwards, the measurements for the starting day (t_0_) were subtracted from later time points (t_3_, t_5_, and t_7_) to correct for initial absorbance of carbon sources and the initial cell density. Once all corrections were done, a growth threshold of 0.08 at OD 595 nm was fixed (other thresholds above and below 0.08 were also tested, but did not alter the results) (Table S5-S7). All values above this threshold were considered as growth, values below the threshold as no growth. Therefore, bacterial growth across the 31 carbon sources was converted into Boolean vectors defining phenotypic profiles. The phenotypic profiles of two partners *P* and *Q* were thus represented by their Boolean vectors of the form *p* = (0, 1,…, 1) and *q* = (0, 1,…, 1), where each vector of size *n* = 31 indicates the number of carbon sources used in this study. Niche width (*N*_*w*_) is given by the number of carbon sources, in which bacteria could grow, and therefore, by the sum of vector entries.

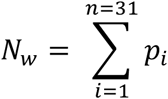

Niche similarity (*N*_*s*_) was simply the intersection between vectors:

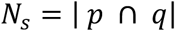

Niche dissimilarity (*N*_*d*_) was determined by subtracting the union from the intersection between vectors (known as *symmetric difference* or *disjunctive union*):

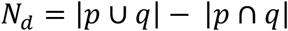

Another way to express the degree of niche dissimilarity between two interacting genotypes is to normalize both i) union minus niche intersection and ii) intersection with the niche sizes occupied by both corresponding partners (i.e., the niche union). To this end, the two measures (i.e., i and ii) were divided by the niche union, thus resulting in the normalized niche dissimilarity (*Jaccard distance*) and the normalized niche similarity (*Jaccard index*), respectively. The Jaccard distance is given by:

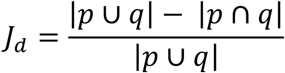

The Jaccard index is defined as:

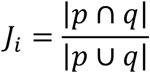

The expected niche (*E*) was calculated as the intersection between the niches utilized by monocultures:

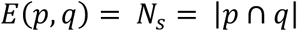

If the observed niche, defined as a Boolean vector with the carbon sources utilized by the coculture is given by *O*, the correctly predicted niche (*N*_*pred*_) is defined as:

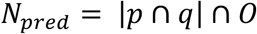

Niche contraction (*N*_*ctr*_) is given by:

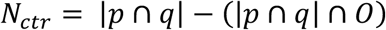

Finally, niche expansion *N*_*ex*_ is defined as:

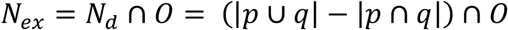

#### Statistical analysis

Statistical differences between groups were inferred by using the non-parametric Mann-Whitney U-test, which does not assume normal distribution of data. A non-parametric Spearman rank correlation test was performed to detect statistical associations between variables. A Gaussian non-linear model was fitted to the dataset (Figs. 3a, b) and measures of goodness of fit between the data and the model were calculated with the function NonlinearModelFit routine as implemented in Mathematica 12 (Table S4).

## Acknowledgements

We thank the entire Kost-lab (present and past) for useful discussion as well as Marita Hermann and Antje Moehlmeyer for technical assistance. Advice on construction of auxotrophic strains by Jenna Gallie (MPI EvoBio) for *Pseudomonas fluorescens* and by Ákos T. Kovács (DTU) for *Bacillus subtilis* is gratefully acknowledged. This work was funded by the German Research Foundation (priority program SPP1617, KO 3909/2-1: CK, SG), (SFB 944, P19: CK), (KO 3909/4-1: CK), (TH 831/3-2: KMT) and the University of Osnabrück (LO).

## Author contributions

Conceptualization: LO, SG, and CK, Experimental design: SG, CK, and NA, Strain construction: SG, MK, and KMT, Performed experiments: NA, SG, Data analysis: LO, Data interpretation: LO, SG, and NA, Writing, LO, SG, NA, and CK, Resources and funding acquisition: CK.

