## Supplementary Information for "Obligate cross-feeding expands the metabolic niche of bacteria"

[#] Authors contributed equally to this work

**Supplementary Figures**

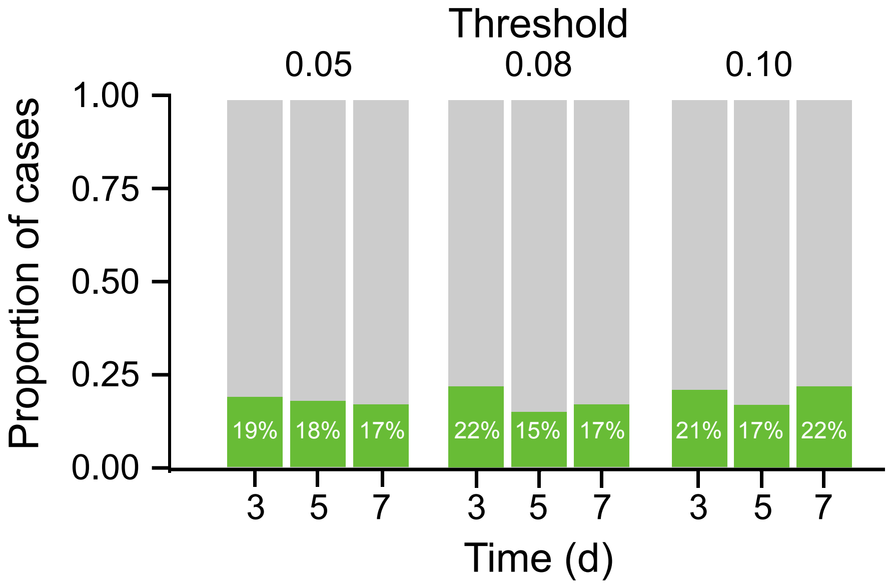

**Fig. S1. Proportion of all cases of niche expansion, in which cocultures could use a certain carbon source, while none of the two monocultures could grow under the same conditions.** Data for the different threshold levels to define growth and different time points of the experiment are shown.

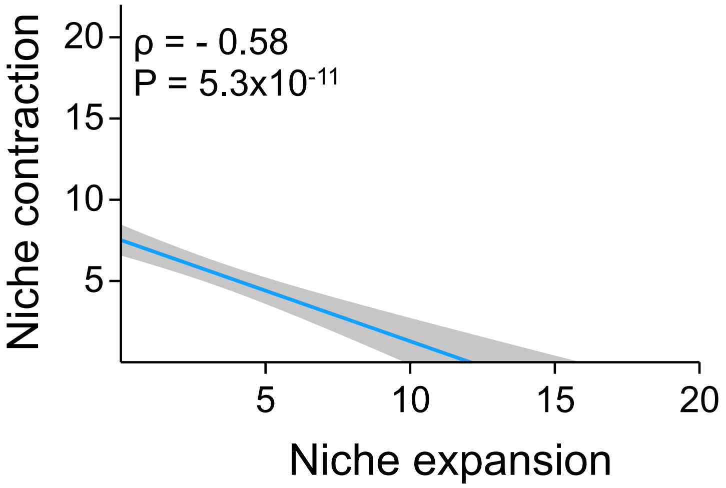

**Fig. S2. Niche contraction is negatively associated with niche expansion.** A linear regression was fitted to the data (blue line, grey area: ±95% confidence interval). The result of a Spearman rank correlation is shown (n = 108).

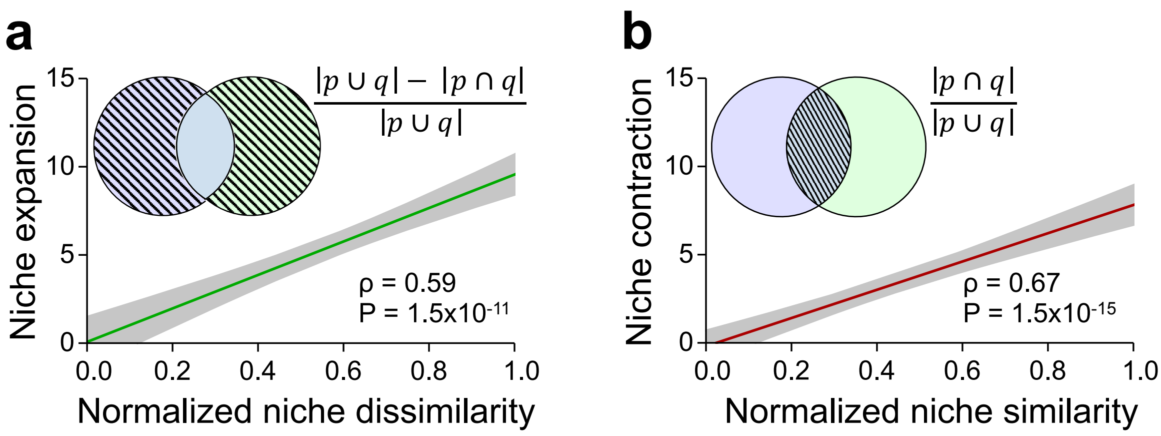

**Fig. S3. Monoculture niches and their normalized overlap can predict the magnitude of niche expansion and niche contraction**. **a** Normalized niche dissimilarity (i.e., Jaccard distance) strongly predicts niche expansion (Spearman rank correlation: $P = 1.5\text{x}10^{-11}$, $\rho = 0.59$, $n = 108$), while **b** normalized niche similarity (i.e., Jaccard index) strongly predicts niche contraction (Spearman rank correlation: $P = 1.5\text{x}10^{-15}$, $\rho = 0.67$, $n = 108$). A linear regression was fitted to the data (green/ red line, grey area: ±95% confidence interval).

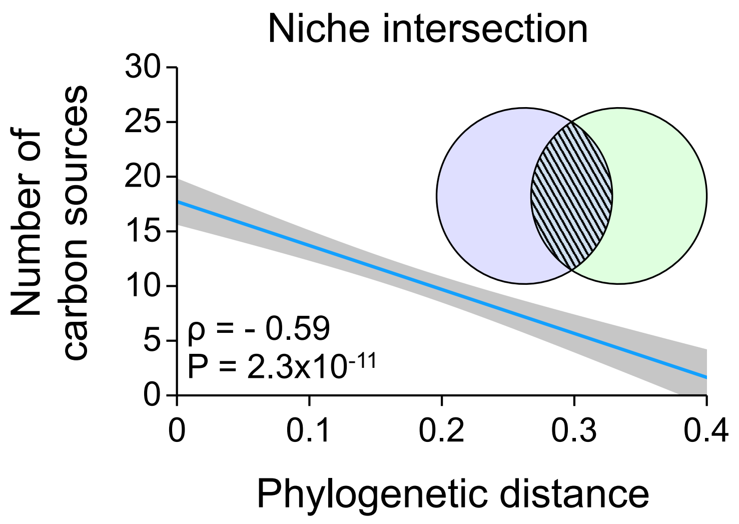

**Fig. S4. Niche intersection is negatively associated with phylogenetic distance.** A linear regression was fitted to the data (blue line, grey area: ±95% confidence interval). The result of a Spearman rank correlation is shown (n = 108).

### Supplementary Tables

#### Table S1. Bacterial strains and plasmid used.

| Strain | Source | Identifier |
|---|---|---|
| *Acinetobacter baylyi* ADP1 | 1 | 2 |
| *Bacillus subtilis* 3610 *comI*$^{Q12L}$ | 3 | Provided by Ákos T. Kovács, DTU (DK1042) |
| *Escherichia coli* BW25113 | 4 | *E. coli* Genetic resources at Yale CGSC, The Coli Genetic Stock Center |
| *Pseudomonas fluorescens* SBW25 | Paul B. Rainey | 5 |
| *Shewanella oneidensis* | Lab stock | Kai Thormann Lab |
| *Escherichia coli* WM3064 | Lab stock | Kai Thormann Lab |
| *Escherichia coli* DH5α λpir | Lab stock | N/A |
| *Acinetobacter baylyi* ADP1*, ΔargH* | This study | N/A |
| *Acinetobacter baylyi* ADP1*, ΔhisD* | This study | N/A |
| *Acinetobacter baylyi* ADP1*, ΔleuB* | This study | N/A |
| *Acinetobacter baylyi* ADP1*, ΔtrpB* | This study | N/A |
| *Bacillus subtilis* 168, *trpC2*, Δ*argH::ermR* | BGSC | BKE29440 |
| *Bacillus subtilis* 168, *trpC2*, Δ*hisD::ermR* | BGSC | BKE34910 |
| *Bacillus subtilis* 168, *trpC2*, Δ*leuB::ermR* | BGSC | BKE28270 |
| *Bacillus subtilis* 3610 *comI*$^{Q12L}$, Δ*argH* | This study | N/A |
| *Bacillus subtilis* 3610 *comI*$^{Q12L}$, Δ*hisD* | This study | N/A |
| *Bacillus subtilis* 3610 *comI*$^{Q12L}$, Δ*leuB* | This study | N/A |
| *Escherichia coli* BW25113*, ΔargH* | This study | N/A |
| *Escherichia coli* BW25113*, ΔhisD* | This study | N/A |
| *Escherichia coli* BW25113*, ΔleuB* | This study | N/A |
| *Escherichia coli* BW25113*, ΔtrpB* | This study | N/A |
| *Pseudomonas fluorescens* SBW25, Δ*leuB* | This study | N/A |
| *Pseudomonas fluorescens* SBW25, Δ*trpB* | This study | N/A |
| *Shewanella oneidensis, ΔargH* | This study | N/A |
| *Shewanella oneidensis, ΔhisD* | This study | N/A |
| *Shewanella oneidensis, ΔleuB* | This study | N/A |
| *Shewanella oneidensis, ΔtrpB* | This study | N/A |

| Plasmids | Source | Identifier |
|---|---|---|
| pCR8/GW/TOPO | Invitrogen, ThermoFisher | Catalog # K250020 |
| pUIC3 | 6 | Lab stock |
| pRK2013 | 7 | Lab stock |
| pCP20 | 8 | *E. coli* Genetic resources at Yale CGSC, The Coli Genetic Stock Center |
| pDR244 | 9 | BGSC |

**Table S2. Chemicals and materials used.**

| Reagent or resource | Source | Identifier |
|---|---|---|
| Lysogeny broth (LB), Lennox | Carl Roth GmbH | Catalog # X964.1 |
| LB agar, Lennox | Carl Roth GmbH | Catalog # X965.3 |
| Dipotassium hydrogen phosphate | Carl Roth GmbH | Catalog # 26931.263 |
| Sodium dihydrogen phosphate | Carl Roth GmbH | Catalog # T879.2 |
| Magnesium sulphate heptahydrate | Carl Roth GmbH | Catalog # P027.2 |
| Potassium chloride | VWR | Catalog # 26764.260 |
| Calcium chloride dihydrate | Carl Roth GmbH | Catalog # 5239.2 |
| Ammonium chloride | VWR | Catalog # 21236.267 |
| Iron (II) sulphate heptahydrate | Merck | Catalog # 3965 |
| Manganese chloride | AppliChem | Catalog # A2087.0100 |
| Cobalt chloride hexahydrate | AppliChem | Catalog # A2087.0100 |
| Boric acid | Carl Roth GmbH | Catalog # 6943.3 |
| Nickel chloride | AppliChem | Catalog # A3917.0100 |
| Zinc sulphate heptahydrate | Carl Roth GmbH | Catalog # K301.1 |
| Copper chloride dihydrate | AppliChem | Catalog # 131264.1210 |
| N-Acetyl-D-glucosamine | Carl Roth GmbH | Catalog # 8993.4 |
| Kanamycin | Carl Roth GmbH | Catalog # T832.2 |
| Erythromycin | Carl Roth GmbH | Catalog # 4166.1 |
| Lincomycin hydrochloride | Sigma- Aldrich | Catalog # 62143-1G |
| D-Cycloserine | Carl Roth GmbH | Catalog # CN37.3 |
| Nitrofurantoin | Sigma- Aldrich | Catalog # N7878 |
| Tetracycline hydrochloride | Carl Roth GmbH | Catalog # 0237.1 |
| Spectinomycin dihydrochloride | Alfa Aesar | Catalog # J61820 |
| 2,6-diaminopimelic acid (DAP) | Alfa Aesar | Catalog # B22391.06 |
| X-gal (5-bromo-4-chloro-3-indolyl-β-D-galactopyranoside) | Carl Roth GmbH | Catalog # 3215.4 |
| Glycerol | Carl Roth GmbH | Catalog # 7533.1 |
| L-Arginine monohydrochloride | AppliChem reagents | Catalog # A3709 |
| L-Histidine monohydrochloride | AppliChem reagents | Catalog # A3733 |
| L-Leucine | AppliChem reagents | Catalog # A3460 |
| L-Tryptophan | AppliChem reagents | Catalog # A3445 |
| 48 well deep well plates | Axygen | Catalog # P-5ML-48-C-S |
| 96 well deep well plates | Eppendorf | Catalog # 0030506308D |
| BiOLOG-Eco-Plate | MediLoc GmbH | Catalog # BLG1506 |
| Plate sealer | Greiner bio-one | Catalog # 676070 |
| Petri dish | Greiner bio-one | Catalog # 633180 |

**Table S3. Primers used.** Primers were used for the construction of auxotrophs of *Pseudomonas fluorescens* (PF) and *Shewanella oneidensis* (SO) as well as to verify the removal of the antibiotic cassette from the genome of *Bacillus subtilis* 3610 *comI*$^{Q12L}$ (BS). UF = upstream forward, UR = upstream reverse, DF = downstream forward, DR = downstream reverse.

| Species | Gene | Amino acid | Primer | Sequence (5'-3') |
|---|---|---|---|---|
| PF | *argH* | Arginine | UF | gaagatct TGAGTCAGGGCAGCAAGC |
| | | | UR | gcagcatgcggatccgttgacggaTGACGGAGGCGGTGAAG |
| | | | DF | tccgtcaacggatccgcatgctgc GTAACCATGTGGGCGGGAC |
| | | | DR | gaagatct TTCGATCAGGGTGGTTTCGG |
| PF | *hisD* | Histidine | UF | gaagatct ACGTTCGCCTGTTGATCGT |
| | | | UR | gcagcatgcggatccgttgacggaCCAGCTCAGCAGATGATCC |
| | | | DF | tccgtcaacggatccgcatgctgc GAGGTGAATCCCTGAGTGCC |
| | | | DR | gaagatct TGGCAACACCTCAAAACCCT |
| PF | *leuB* | Leucine | UF | gaagatctGATCAATGTGGGAGCTGGCT |
| | | | UR | gcagcatgcggatccgttgacggaGAGCTCCAGTACCTTGACCG |
| | | | DF | tccgtcaacggatccgcatgctgc ACATCTGGTCGCAGGGTTG |
| | | | DR | gaagatct GGCCGAGGATCTTGTTGGTC |
| PF | *trpB* | Tryptophan | UF | gaagatctCAGGTCCTCTGCCACCAATG |
| | | | UR | gcagcatgcggatccgttgacggaAGGGTTTCAGCGACGTAACG |
| | | | DF | tccgtcaacggatccgcatgctgc CGTGGCGACAAAGACATGC |
| | | | DR | gaagatct GTCTTGCTGTTTAGGCCGTG |
| SO | *argH* | Arginine | UF | gcgaattcgtggatccagATCCGAAAACCGTTTAGTGGGC |
| | | | UR | tcacttcgccattaacccgcCCATAAAGCCATGCTCAATCC |
| | | | DF | gattgagcatggctttatggGCGGGTTAATGGCGAAGTGA |
| | | | DR | gccaagcttctctgcaggatCGCCATGATATTGCAGGTAAA |
| SO | *hisD* | Histidine | UF | gcgaattcgtggatccagatCCCAACCTGCTGCGTCGTTT |
| | | | UR | acggcttaagtcgttcagccCATATCCATAGATCACCCCATCATTTT |
| | | | DF | tggggtgatctatggatatgGGCTGAACGACTTAAGCCGT |
| | | | DR | gccaagcttctctgcaggatGGCGTACATGCCGTATGTCG |
| SO | *leuB* | Leucine | UF | gcgaattcgtggatccagatCGACTTAGAAGCCTTAGCTTTTAT AGAGGC |
| | | | UR | aggaactgtcatcacacaccCATACGCCCGCCCACTCCTT |
| | | | DF | aaggagtgggcgggcgtatgGGTGTGTGATGACAGTTCCTT |
| | | | DR | gccaagcttctctgcaggatGCAAAGTTTGGGTTGCCAAT |

| | | | | |
|---|---|---|---|---|
| SO | *trpB* | Tryptophan | UF | gcgaattcgtggatccagaTGCAGTGAAACCGAGTTAGAA |
| | | | UR | gttcatcttagttattcCTCCTGTGACATACCTTGTTCCT |
| | | | DF | aggaacaaggtatgtcacagGAGGAATAACTAAGATGAACAGCAT TTCAAACC |
| | | | DR | gccaagcttctctgcaggatGGCGCAATGCCATGGGCTTT |
| BS | *argH* | Arginine | UF | GCTTCAGGAGCAAGGATATAATG |
| | | | DR | CACAAGGTCTGAGTTCTCCTCAC |
| BS | *hisD* | Histidine | UF | CAGCCACATGAGCTATTACACAG |
| | | | DR | GAAGTAAGCGTAGCCTTGTTCTG |
| BS | *leuB* | Leucine | UF | GCCAGTGTTATCAATCTTCCG |
| | | | DR | AGGAACCGATAAATACGTGCTC |

**Table S4. Software used.**

| Software | Source | Identifier |
|---|---|---|
| MEGA X | 10 | https://www.megasoftware.net/ |
| Primer3 | 11 | http://biotools.umassmed.edu/bioapps/ primer3_www.cgi |
| iTOL | 12 | http://itol.embl.de/ |
| Softmax Pro 6 software | Molecular Devices, San Jose, CA, USA | N/A |
| IBM SPSS statistics 26 | IBM Corp. Released 2019. IBM SPSS Statistics | https://www.ibm.com/support/pages/downloading-ibm-spss-statistics-26 |
| Mathematica | N/A | https://www.wolfram.com/mathematica/ |

**Table S5. Analysing different time points and decreasing the threshold limit to define growth (i.e., $OD_{595nm} \geq 0.05$) does not change the main conclusions.**

| Time point | Statistical comparison | P-value | Statistics | n | Figure |
|---|---|---|---|---|---|
| 3 days | Predicted-expansion | 0.1 | U= 6572 | 108 | Fig. 2 |
| | Predicted-contraction | $2.5x10^{-10}$ | U= 8721.5 | 108 | |
| | Expansion-contraction | $4.9x10^{-8}$ | U= 3341 | 108 | |
| | Niche expansion 3D | $10^{-4}$ | R= 0.7 | 108 | Fig. 3 |
| | Niche contraction 3D | $10^{-4}$ | R= 0.75 | 108 | |
| | Niche expansion-(union-intersection) | $1.9x10^{-8}$ | ρ= 0.51 | 108 | |
| | Niche contraction-intersection | $2.1x10^{-15}$ | ρ= 0.67 | 108 | |
| | Niche expansion-phylogenetic distance | $2.3x10^{-3}$ | ρ= 0.29 | 108 | Fig. 4 |
| | Niche contraction-phylogenetic distance | $4.1x10^{-3}$ | ρ= - 0.27 | 108 | |
| | Niche expansion-(within-between species) | 0.037 | U= 672.5 | 108 | |
| | Niche contraction-(within-between species) | 0.88 | U= 965.5 | 108 | |
| | Niche intersection-phylogenetic distance | $4.9x10^{-10}$ | ρ= - 0.55 | 108 | Fig. S1 |
| | Niche contraction-niche expansion | $6.7x10^{-11}$ | ρ= -0.58 | 108 | Fig. S2 |
| 5 days | Predicted-expansion | 0.02 | U= 6940.5 | 108 | Fig. 2 |
| | Predicted-contraction | $1.0x10^{-11}$ | U= 8942.5 | 108 | |
| | Expansion-contraction | $4.6x10^{-8}$ | U= 3335 | 108 | |
| | Niche expansion 3D | $10^{-4}$ | R= 0.75 | 108 | Fig. 3 |
| | Niche contraction 3D | $10^{-4}$ | R= 0.74 | 108 | |
| | Niche expansion-(union-intersection) | $6.5x10^{-14}$ | ρ= 0.64 | 108 | |
| | Niche contraction-intersection | $3.2x10^{-18}$ | ρ= 0.72 | 108 | |
| | Niche expansion-phylogenetic distance | $7.5x10^{-9}$ | ρ= 0.52 | 108 | Fig. 4 |
| | Niche contraction-phylogenetic distance | $7.6x10^{-6}$ | ρ= -0.42 | 108 | |
| | Niche expansion-(within-between species) | $5.9x10^{-4}$ | U= 496 | 108 | |
| | Niche contraction-(within-between species) | 0.18 | U= 1119.5 | 108 | |
| | Niche intersection-phylogenetic distance | $9.2x10^{-11}$ | ρ= - 0.57 | 108 | Fig. S2 |
| | Niche contraction-niche expansion | $1.3x10^{-12}$ | ρ= - 0.62 | 108 | Fig.S3 |
| 7 days | Predicted-expansion | 0.02 | U= 6874.5 | 108 | Fig. 2 |
| | Predicted-contraction | $2.9x10^{-14}$ | U= 9298.5 | 108 | |

| | | | | |
|---|---|---|---|---|
| Expansion-contraction | 6.5x10$^{-16}$ | U= 2142.5 | 108 | |
| Niche expansion 3D | 10$^{-4}$ | R= 0.77 | 108 | Fig. 3 |
| Niche contraction 3D | 10$^{-4}$ | R= 0.71 | 108 | |
| Niche expansion-(union-intersection) | 3.8x10$^{-13}$ | ρ= 0.63 | 108 | |
| Niche contraction-intersection | 1.1x10$^{-17}$ | ρ= 0.71 | 108 | |
| Niche expansion-phylogenetic distance | 4.8x10$^{-8}$ | ρ= 0.50 | 108 | Fig. 4 |
| Niche contraction-phylogenetic distance | 5.8x10$^{-8}$ | ρ= - 0.49 | 108 | |
| Niche expansion-(within-between species) | 3.9x10$^{-4}$ | U= 481.5 | 108 | |
| Niche contraction-(within-between species) | 0.056 | U= 1191 | 108 | |
| Niche intersection-phylogenetic distance | 4.1x10$^{-10}$ | ρ= - 0.56 | 108 | Fig. S2 |
| Niche contraction-niche expansion | 9.8x10$^{-13}$ | ρ= -0.62 | 108 | Fig. S3 |

**Table S6. Analysing different time points does not change the main conclusions.** The threshold limit to define growth was $OD_{595nm} \geq 0.08$. Grey-shaded area (i.e. 5 days) represents the time point used for figures in the main text.

| Time point | Statistical comparison | P-value | Statistics | n | Figure |
|---|---|---|---|---|---|
| 3 days | Predicted-expansion | 0.25 | U= 6358.5 | 108 | Fig. 2 |
| | Predicted-contraction | 3.6x10$^{-5}$ | U= 7716 | 108 | |
| | Expansion-contraction | 7.7x10$^{-5}$ | U= 4029.5 | 108 | |
| | Niche expansion 3D | 10$^{-4}$ | R= 0.65 | 108 | Fig. 3 |
| | Niche contraction 3D | 10$^{-4}$ | R= 0.78 | 108 | |
| | Niche expansion-(union-intersection) | 4.7x10$^{-5}$ | ρ= 0.38 | 108 | |
| | Niche contraction-intersection | 1.2x10$^{-19}$ | ρ= 0.74 | 108 | |
| | Niche expansion-phylogenetic distance | 4.8x10$^{-4}$ | ρ= 0.33 | 108 | Fig. 4 |
| | Niche contraction-phylogenetic distance | 4.0x10$^{-5}$ | ρ= -0.38 | 108 | |
| | Niche expansion-(within-between species) | 0.037 | U= 672.5 | 108 | |
| | Niche contraction-(within-between species) | 0.66 | U= 1002 | 108 | |
| | Niche intersection-phylogenetic distance | 4.9x10$^{-10}$ | ρ= -0.55 | 108 | Fig. S2 |
| | Niche contraction-niche expansion | 6.7x10$^{-11}$ | ρ= -0.58 | 108 | Fig. S3 |
| 5 days | Predicted-expansion | 0.02 | U= 6866.5 | 108 | Fig. 2 |
| | Predicted-contraction | 8.1x10$^{-8}$ | U= 8277.5 | 108 | |
| | Expansion-contraction | 7x10$^{-5}$ | U= 4020.5 | 108 | |
| | Niche expansion 3D | 10$^{-4}$ | R= 0.65 | 108 | Fig. 3 |
| | Niche contraction 3D | 10$^{-4}$ | R= 0.72 | 108 | |
| | Niche expansion-(union-intersection) | 4.03x10$^{-9}$ | ρ= 0.53 | 108 | |
| | Niche contraction-intersection | 1.16x10$^{-19}$ | ρ= 0.74 | 108 | |
| | Niche expansion-phylogenetic distance | 5.8x10$^{-8}$ | ρ= 0.493 | 108 | Fig. 4 |
| | Niche contraction-phylogenetic distance | 1.3x10$^{-8}$ | ρ= - 0.51 | 108 | |
| | Niche expansion-(within-between species) | 6.0x10$^{-5}$ | U= 421.5 | 108 | |
| | Niche contraction-(within-between species) | 0.047 | U= 1202 | 108 | |
| | Niche intersection-phylogenetic distance | 2.3x10$^{-11}$ | ρ= - 0.59 | 108 | Fig. S2 |
| | Niche contraction-niche expansion | 5.3x10$^{-11}$ | ρ= - 0.58 | 108 | Fig. S3 |
| 7 days | Predicted-expansion | 0.02 | U= 6890.5 | 108 | Fig. 2 |

| | | | | |
|---|---|---|---|---|
| Predicted-contraction | $5.7 \times 10^{-10}$ | U= 8659 | 108 | |
| Expansion-contraction | $1.6 \times 10^{-7}$ | U= 3443 | 108 | |
| Niche expansion 3D | $10^{-4}$ | R= 0.67 | 108 | Fig. 3 |
| Niche contraction 3D | $10^{-4}$ | R= 0.74 | 108 | |
| Niche expansion-(union-intersection) | $8.24 \times 10^{-13}$ | ρ= 0.62 | 108 | |
| Niche contraction-intersection | $7.0 \times 10^{-20}$ | ρ= 0.74 | 108 | |
| Niche expansion-phylogenetic distance | $1.54 \times 10^{-8}$ | ρ= 0.51 | 108 | Fig. 4 |
| Niche contraction-phylogenetic distance | $4.0 \times 10^{-10}$ | ρ= - 0.56 | 108 | |
| Niche expansion-(within-between species) | $2.9 \times 10^{-6}$ | U= 334 | 108 | |
| Niche contraction-(within-between species) | 0.05 | U= 1194.5 | 108 | |
| Niche intersection-phylogenetic distance | $1.4 \times 10^{-10}$ | ρ= - 0.57 | 108 | Fig. S2 |
| Niche contraction-niche expansion | $2.0 \times 10^{-7}$ | ρ= -0.47 | 108 | Fig. S3 |

**Table S7. Analysing different time points and increasing the threshold limit to define growth (i.e., $OD_{595nm}$ ≥ 0.1) does not change the main conclusions.**

| Time point | Statistical comparison | P-value | Statistics | n | Figure |
|---|---|---|---|---|---|
| 3 days | Predicted-expansion | 0.42 | U= 6198 | 108 | Fig. 2 |
| | Predicted-contraction | 1.7x$10^{-3}$ | U= 7255 | 108 | |
| | Expansion-contraction | 4.5x$10^{-3}$ | U= 4538.5 | 108 | |
| | Niche expansion 3D | $10^{-4}$ | R= 0.64 | 108 | Fig. 3 |
| | Niche contraction 3D | $10^{-4}$ | R= 0.79 | 108 | |
| | Niche expansion-(union-intersection) | 1.9x$10^{-4}$ | ρ= 0.35 | 108 | |
| | Niche contraction-intersection | 2.9x$10^{-22}$ | ρ= 0.77 | 108 | |
| | Niche expansion-phylogenetic distance | 4.0x$10^{-4}$ | ρ= 0.33 | 108 | Fig. 4 |
| | Niche contraction-phylogenetic distance | 7.7x$10^{-6}$ | ρ= - 0.42 | 108 | |
| | Niche expansion-(within-between species) | 0.012 | U= 619.5 | 108 | |
| | Niche contraction-(within-between species) | 0.46 | U= 1040 | 108 | |
| | Niche intersection-phylogenetic distance | 4.9x$10^{-10}$ | ρ= - 0.55 | 108 | Fig. S1 |
| | Niche contraction-niche expansion | 6.7x$10^{-11}$ | ρ= - 0.58 | 108 | Fig. S2 |
| 5 days | Predicted-expansion | 0.03 | U= 6839.5 | 108 | Fig. 2 |
| | Predicted-contraction | 7.9x$10^{-7}$ | U= 8076 | 108 | |
| | Expansion-contraction | 4.3x$10^{-4}$ | U= 4234 | 108 | |
| | Niche expansion 3D | $10^{-4}$ | R= 0.63 | 108 | Fig. 3 |
| | Niche contraction 3D | $10^{-4}$ | R= 0.70 | 108 | |
| | Niche expansion-(union-intersection) | 8.8x$10^{-8}$ | ρ= 0.49 | 108 | |
| | Niche contraction-intersection | 1.0x$10^{-18}$ | ρ= 0.72 | 108 | |
| | Niche expansion-phylogenetic distance | 3.0x$10^{-8}$ | ρ= 0.5 | 108 | Fig. 4 |
| | Niche contraction-phylogenetic distance | 9.9x$10^{-9}$ | ρ= -0.51 | 108 | |
| | Niche expansion-(within-between species) | 1.4x$10^{-5}$ | U= 379 | 108 | |
| | Niche contraction-(within-between species) | 0.046 | U= 1201.5 | 108 | |
| | Niche intersection-phylogenetic distance | 5.9x$10^{-11}$ | ρ= - 0.58 | 108 | Fig. S2 |

| | | | | | |
|---|---|---|---|---|---|
| | Niche contraction-niche expansion | $3.3 \times 10^{-7}$ | ρ= - 0.47 | 108 | Fig. S3 |
| 7 days | Predicted-expansion | 0.04 | U= 6758.5 | 108 | Fig. 2 |
| | Predicted-contraction | $2.9 \times 10^{-10}$ | U= 8696 | 108 | |
| | Expansion-contraction | $3.6 \times 10^{-8}$ | U= 3326 | 108 | |
| | Niche expansion 3D | $10^{-4}$ | R= 0.62 | 108 | Fig. 3 |
| | Niche contraction 3D | $10^{-4}$ | R= 0.67 | 108 | |
| | Niche expansion-(union-intersection) | $5.8 \times 10^{-10}$ | ρ= 0.55 | 108 | |
| | Niche contraction-intersection | $5.3 \times 10^{-14}$ | ρ= 0.64 | 108 | |
| | Niche expansion-phylogenetic distance | $1.1 \times 10^{-6}$ | ρ= 0.45 | 108 | Fig. 4 |
| | Niche contraction-phylogenetic distance | $9.6 \times 10^{-8}$ | ρ= - 0.49 | 108 | |
| | Niche expansion-(within-between species) | $7.2 \times 10^{-6}$ | U= 360 | 108 | |
| | Niche contraction-(within-between species) | 0.1 | U= 1149.5 | 108 | |
| | Niche intersection-phylogenetic distance | $1.5 \times 10^{-10}$ | ρ= - 0.57 | 108 | Fig. S2 |
| | Niche contraction-niche expansion | $3.9 \times 10^{-5}$ | ρ= -0.38 | 108 | Fig. S3 |
